# Coding Agents as a Mechanism for Formalizing and Transferring Domain Knowledge in DNA Origami Design

**DOI:** 10.64898/2026.04.11.717962

**Authors:** Daniel Fu, Yonggang Ke

## Abstract

This work investigates the use of a coding agent to design and manipulate DNA origami structures through the caDNAno2 design interface. The design methodology produced by the agent is saved as transferable, repeatable text-file instructions interpretable by another coding agent, converting informal domain expertise into formalized, distributable protocols. We demonstrate this through a parametric pipeline for simple single and multi-layer designs, identifying several failure modes that each required explicit domain knowledge from the human designer. Once encoded, this knowledge enabled the agent to autonomously extend designs to arbitrary lattice dimensions, generate structures to physical specifications, execute targeted edits from natural language prompts, and run the full pipeline from design through molecular dynamics simulation.

## 1. Introduction

Modifying a DNA origami design requires re-routing the scaffold, re-placing staples, and re-verifying the structure for each parameter change^1,2^. A single redesign of a simple structure may be quick, taking several minutes, but multiple iterations or greater complexity accumulate to hours of repetitive, error-prone work. The knowledge required to correctly perform these tasks is well documented for human practitioners^1^, but is not encoded in the design tools themselves. The tools implement the operations, but the rules governing when and how to compose those operations correctly remain in tutorials, lab conventions, and individual expertise. A coding agent can bridge this gap. It can read the tools’ source code to learn what operations are available, acquire the composition rules through interaction with a designer, and produce executable artifacts that encode both together. The result is that a designer’s expertise, once transferred, becomes a reproducible workflow that any subsequent user or agent can apply without re-learning the rules from scratch.

The effort to algorithmically accelerate the design process and eliminate design tedium has been approached from various angles. A substantial ecosystem of automated design tools now exists, spanning lattice-based design^3^, automated staple breaking^2^, curved and tubular structures^4^, wireframe origami from 2D and 3D meshes^5–8^, shape annealing^9^, and interactive 3D editing environments^10^. However, there is little continuity between these tools. Each addresses a specific shape class or design subtask, embodying its own design principles and operating through a disconnected interface. In practice, the adoption friction of learning each new tool means that many designers default to a single familiar environment, even when a more specialized tool exists for their task. A coding agent that can read any open-source tool’s source code could serve as a natural language interface across all of them, allowing designers to adopt new methodologies without learning each tool’s interface individually. In this work, we initiate that investigation by retracing the DNA origami design chronology^11–15^, from the beginning; folding simple single and multi-layer designs, and developing the methodology for an agent-mediated pipeline that consolidates design principles, techniques, and tools toward greater structural complexity.

A coding agent is a large language model (LLM) that operates by reading files, writing and executing code, and iterating on the results within a command-line environment. Unlike chatbot-style LLMs that produce only text responses, coding agents take actions. They can navigate a codebase, call APIs, run scripts, inspect outputs, and modify files across multiple steps without human intervention between steps. Recent advances in LLM reasoning and tool use have made these agents capable of sustained, multi-step software engineering tasks^16,17^, including navigating unfamiliar codebases, integrating independently developed software packages, and writing domain-specific scripts from documentation and source code alone. However, LLMs have no geometric understanding. They cannot reason about nucleotide positions in 3D space. caDNAno provides a layer of abstraction that makes this unnecessary. It encodes helix connectivity, crossover placement, and strand routing as structured data rather than spatial coordinates, reducing DNA origami design to operations on lists and indices. This abstraction makes caDNAno a tractable conduit for investigating whether a coding agent can acquire the domain knowledge needed to manipulate DNA origami designs through direct interaction with the software, rather than with the 3D structure itself.

In the current work, we demonstrate how a coding agent with direct access to caDNAno’s source code can read internal data models, write and execute Python scripts, and integrate external tools, like tacoxDNA^18^ and oxDNA^19–22^, into an end-to-end pipeline. Critically, the methodology that emerges from this process is not ephemeral. It is saved as verifier functions, parametric scripts, and documented failure catalogs, all of which are transferable, repeatable text-file instructions interpretable by another coding agent. This converts *ad hoc* design tasks into repeatable, iterable procedures and provides a mechanism for converting established design principles^1^ into machine-executable code.

As a proof of concept, we demonstrate the construction of a parametric pipeline for a 2-layer rectangular origami with a tunable cavity using a coding agent (Claude Code). Achieving functional usage required iterating through several distinct failure modes, each corresponding to domain knowledge invisible in the source code (e.g., honeycomb grid-to-layer mapping, crossover parity rules, cavity orientation conventions). Each failure was resolved by explicit correction from the human designer, after which the agent did not repeat that error class. Once this knowledge was encoded, the agent autonomously extended the base template to arbitrary lattice dimensions, generated designs to physical specifications, executed targeted structural edits from natural language prompts, and ran the full pipeline from design through stapling to oxDNA molecular dynamics simulation.

## 2. Tool-Calling LLM: Insufficient Domain Knowledge from Schemas

Before describing the established pipeline, we distinguish two architectures for applying an LLM to a software tool. In tool-calling, the model selects from predefined functions. In code generation, the model reads source code and writes scripts. This section and the next describe our experience with each. The distinction determines what kinds of domain knowledge the model can acquire and apply.

Our initial approach was to embed an LLM directly into the caDNAno GUI as a tool-calling agent (Figure 1, A): the user types a natural language command, the model parses the intent, and calls predefined Python methods that modify the design. The premise was that if we could decompose DNA origami manipulation into a sufficient set of primitives (identify crossovers, move them, place strands, break staples), the model would be able to compose these primitives to execute arbitrary design instructions.

**Figure 1:**
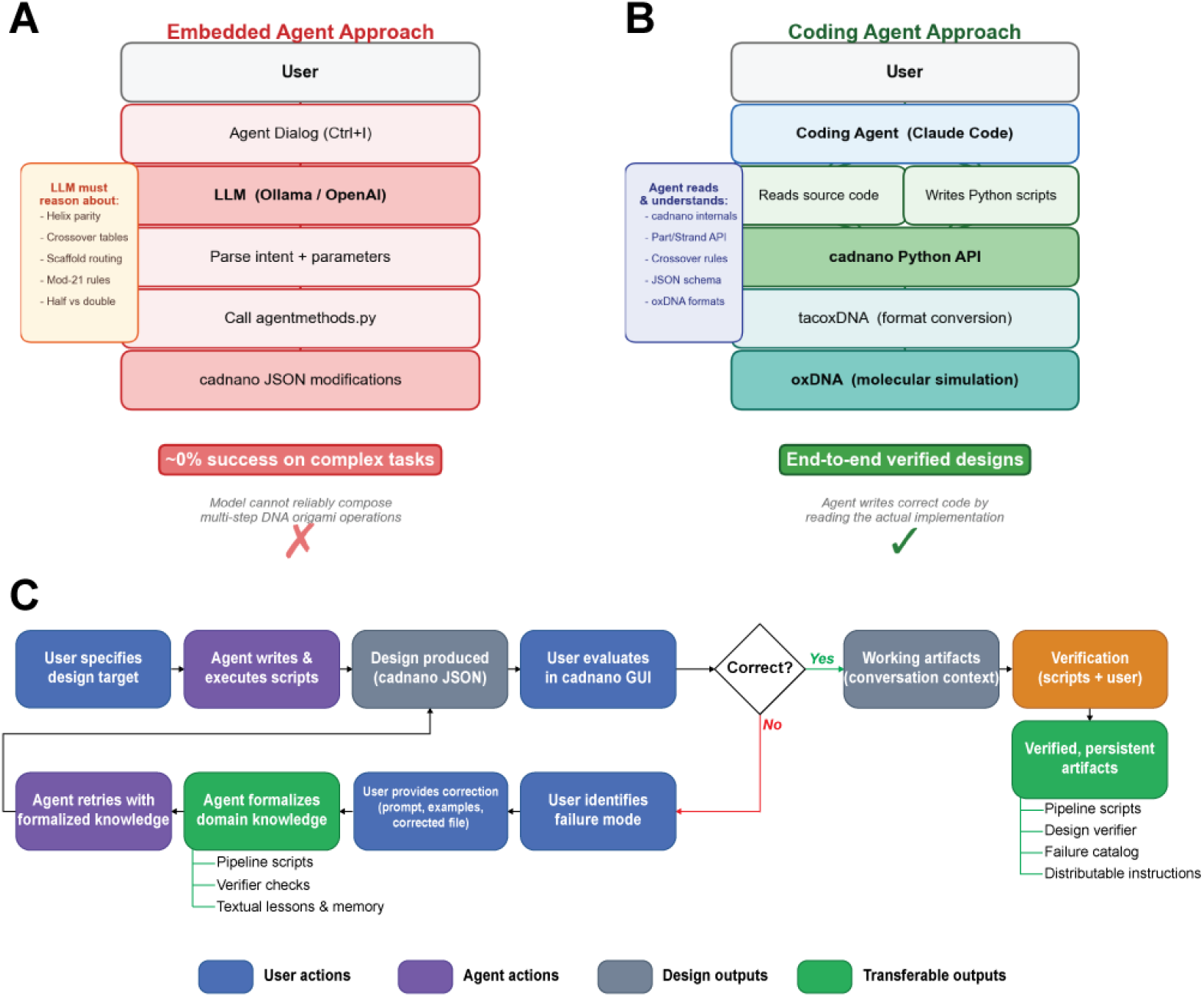
Architecture comparison. (A) Embedded tool-using LLM achieved 0% success on multi-step design tasks because the model could not resolve contextual constraints (helix parity, crossover directionality) from tool schemas alone. (B) Coding agent with source code access produced end-to-end verified designs by reading caDNAno internals, writing Python scripts, and integrating external simulation tools. (C) Knowledge formalization workflow. The user specifies a design target; the agent writes and executes scripts; the user evaluates the result. On failure, the user identifies the failure mode and provides a correction, which the agent formalizes as pipeline scripts, verifier checks, and persistent lessons. The agent retries with this formalized knowledge until the design is correct. On success, working artifacts pass through verification to become persistent, distributable outputs: parametric pipelines, an automated design verifier, a failure catalog, and text-file instructions interpretable by any agent.

In practice, each primitive required contextual parameters that the model could not resolve from the tool schema. A “place crossover” function must distinguish half-crossovers from full crossovers, left-edge from right-edge from interior positions, and 5′→ 3′ from 3′→ 5′ scaffold direction on the target helix. Attempts to encode these distinctions into the API surface produced a combinatorial expansion of tool variants without improving task success. Multi-step instructions such as scaffold routing (placing crossovers to form one continuous scaffold loop) achieved 0% success across GPT-class models. The model could invoke individual tools but could not satisfy the inter-step constraints that govern valid compositions. Reinforcement learning with a design verifier as reward signal^23^ also failed. A local model (Qwen 1.7B) could not discover correct action sequences through exploration, which is unsurprising given that substantially more capable GPT-class models achieved 0% success on the same tasks. The GPT-class failure also precluded distillation, as no successful trajectories were available to serve as training data for the local model. Human-demonstrated trajectories were not evaluated either, as the volume of demonstrations required to adequately cover the constraint space remains an open question.

The limitation appeared to be architectural. A predefined tool API encodes necessary but insufficient domain knowledge. The model can see the available actions but not the reasoning that governs when and how to compose them. Wang et al. demonstrated that tool-calling architectures are systematically inferior to code-generating agents for complex multi-step tasks, precisely because code provides arbitrary composition through control flow, state management, and dynamic parameter construction, capabilities that fixed function schemas cannot express^24^. Where a tool-calling agent must select from a predetermined menu of operations, a coding agent can read source code to understand *why* an operation exists, discover undocumented methods, and compose novel sequences that no API designer anticipated.

## 3. Code-Generating Agent: Domain Knowledge from Source Code

We thus replaced the tool-calling architecture with a coding agent (Claude Code) that reads caDNAno’s source code directly and writes Python scripts (Figure 1, B). Source code access provided two capabilities absent from the tool-calling approach.

- **Discovery through code reading**. The agent identified internal methods not exposed through any public API by reading the strand model source. For example, when the public `createXover` method broke scaffold continuity by splitting strands, the agent located `setConnection3p/5p` in the strand model internals, which preserves connectivity.
- **End-to-end pipeline construction**. The agent integrated caDNAno with tacoxDNA (format conversion) and oxDNA (molecular dynamics) into a single automated pipeline. Each tool is open-source and individually straightforward, but the manual workflow is disjointed, requiring the user to navigate between separate interfaces, file formats, and documentation. The agent automated this by reading each tool’s source code, resolving format requirements, and writing wrapper scripts.

However, source code access alone was not sufficient. Building a correct DNA origami design required domain knowledge that is not encoded in the codebase, such as lattice geometry conventions, crossover placement rules, and scaffold routing patterns. Acquiring this knowledge required iterative human correction. The following sections describe categories of domain knowledge the agent could not derive from code, the process by which each was resolved, and the artifacts that formalized each correction.

## 4. Formalizing Helix Placement on the Honeycomb Lattice

During the iterative correction process, the agent could produce outputs with incorrect lattice dimensions, fragmented scaffold routing, and disconnected strands (Figure 2, A). A cold-start agent given only the instruction to build a 2-layer rectangle can produce a reasonable first attempt, but as additional domain concepts were introduced (half-crossovers, full crossovers, midseam patterns), the agent’s understanding appeared to drift before stabilizing through formalization. The resolution in each case was the same. The user provided an explicit correction, and the agent encoded the rule as a persistent constraint.

**Figure 2:**
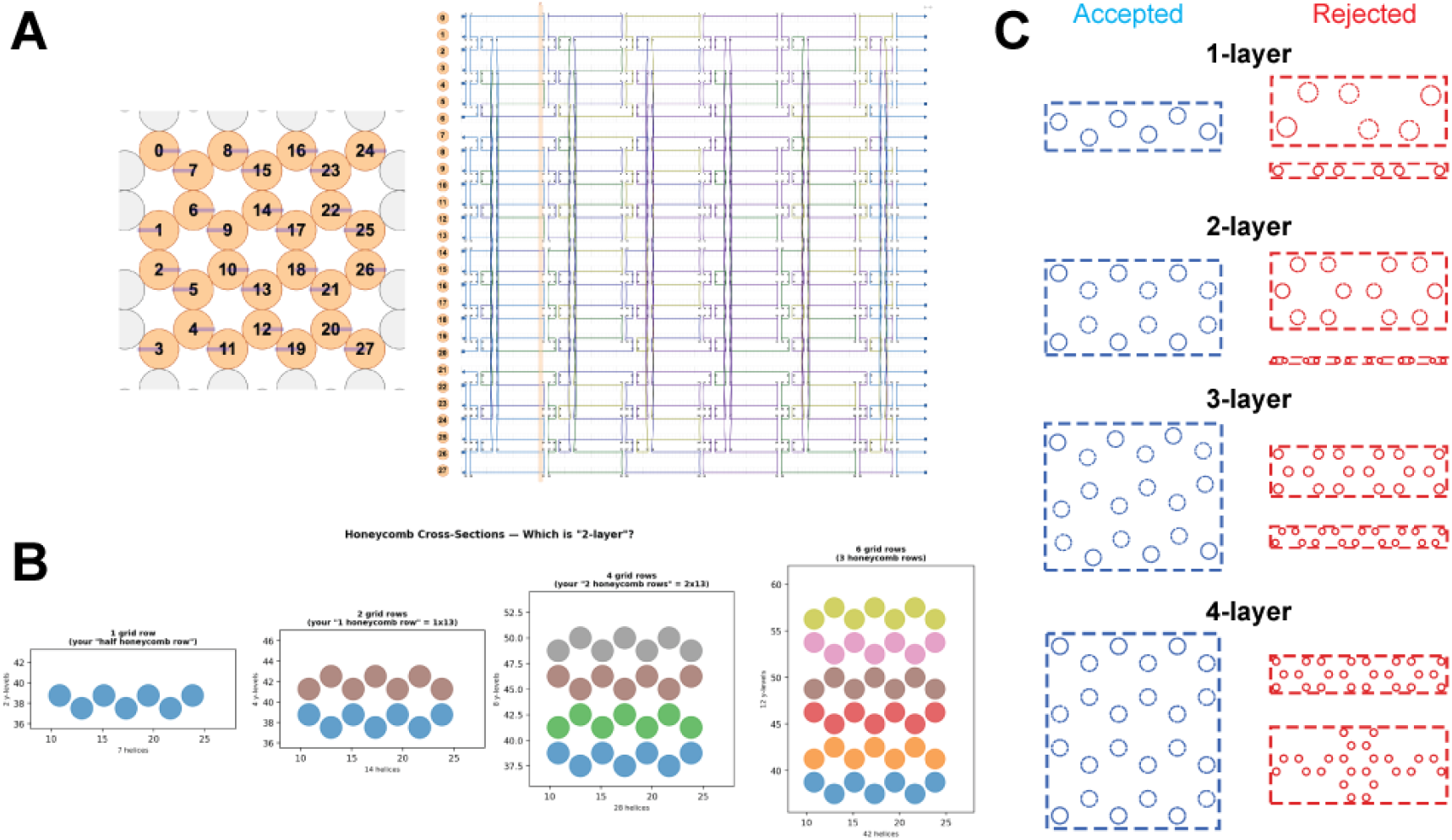
Formalizing helix placement on the honeycomb lattice. (A) Early agent output: a 4-by-7 grid, the agent’s incorrect interpretation of “2-layer,” producing 28 helices with fragmented scaffold routing. (B) Multiple-choice diagram the agent generated to resolve the “layer” ambiguity. When instructed to build a “2-layer” structure, the agent could not determine the mapping between grid rows and physical layers from the source code, and prompted the user to identify the correct correspondence. The user labeled the 2-grid-row option, and the agent retained this for all subsequent designs. (C) With the layer definition resolved, the agent independently developed geometric verification by converting candidate designs to 3D coordinates via tacoxDNA and computing PCA cross-sections. This allowed the agent to distinguish correct flat sheets (aspect ratio ≈ 0.05) from incorrect grid arrangements (≈ 0.84) without human feedback.

### 4.1 The “Layer” Ambiguity

One obstacle that arose during the correction process was the relationship between grid rows and physical layers on the honeycomb lattice. Helices are arranged in a hexagonal packing where adjacent rows are offset vertically. A single row of hexagonally packed helices occupies two distinct y-coordinates, because the vertices of a hexagon do not lie on a single horizontal line. During iterative teaching, the agent at one point constructed a 4-row structure when instructed to build a “2-layer” rectangle, interpreting the two y-coordinates as two physical layers. A 2-layer structure actually corresponds to 2 grid rows on the honeycomb lattice, because one physical layer comprises helices at both y-coordinates of a single hexagonal row.

This ambiguity could not be resolved from the source code. The agent independently generated a multiple-choice diagram of 1-layer, 2-layer, 3-layer, and 4-layer cross-sections and asked the user to identify which arrangement corresponded to “2-layer” (Figure 2, B). The user labeled the correct option, and the agent did not repeat this error in any subsequent session. The correction was formalized as a constraint in the pipeline, where N-layer corresponds to N grid rows on the honeycomb lattice.

### 4.2 Autonomous Geometric Verification via tacoxDNA

With the layer definition resolved, the agent independently developed geometric verification through the end-to-end pipeline (Figure 2, C). caDNAno’s 2D strand routing view provides no spatial information about the 3D conformation of a design. However, with tacoxDNA integrated, the agent could convert candidate designs to 3D coordinates and analyze the resulting geometry. When prompted to verify that a design was physically “flat,” the agent independently devised a verification pipeline. It generated candidate grid layouts, converted each to oxDNA format, computed cross-sectional slices by PCA decomposition of the 3D nucleotide coordinates, and evaluated whether the resulting profile matched a flat rectangle. By measuring cross-sectional circularity (flat sheet: aspect ratio ≈ 0.05; incorrect 2-by-3 grid ≈ 0.84), the agent could reject wrong helix arrangements without human feedback. This behavior was entirely unprompted. The agent was asked to verify flatness, not told how to do so, and independently identified the 3D conversion tools as a means to answer a question that the 2D design interface could not.

## 5. Formalizing Scaffold Routing Rules

With lattice geometry resolved, the next class of errors concerned scaffold routing, specifically how crossovers connect the scaffold strand into a single continuous loop across all helices.

### 5.1 Early Attempts

The agent’s first task was to understand the width of the strand routing design space, specifically how many helices, how long, and where strands begin and end (Figure 3, A, top). After the user introduced the concepts of full crossovers (double crossovers connecting adjacent helices in the interior) and half crossovers (single crossovers at the edges serving as scaffold turns), the agent attempted to place them (Figure 3, A, bottom). However, without knowledge of where these crossovers should go, the agent produced disconnected scaffold fragments rather than a single continuous strand.

**Figure 3:**
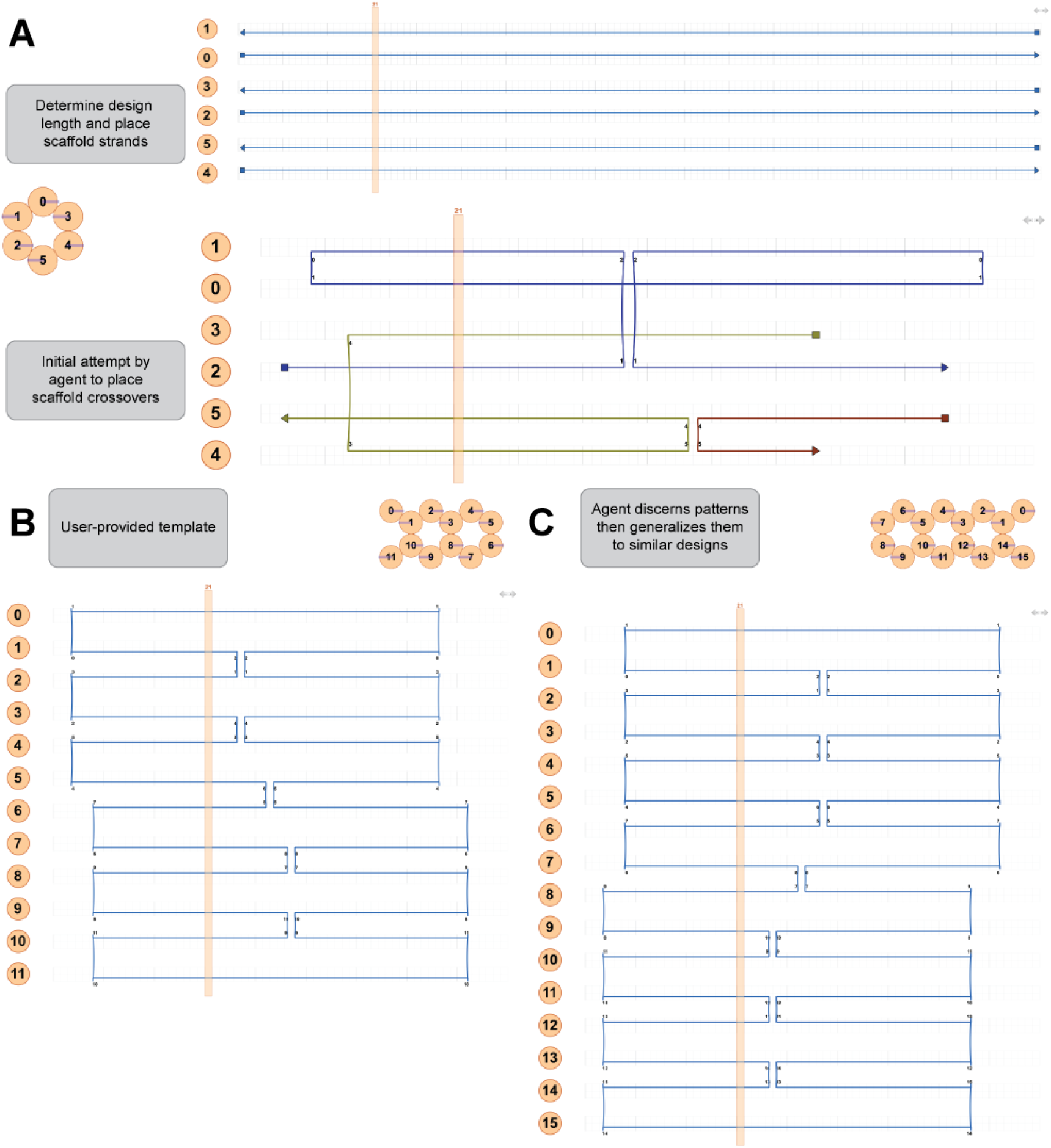
Formalizing scaffold routing. (A) Early agent attempts. Top: the agent explores the path design space, placing strands at various lengths across helices. Bottom: after being told about full and half crossovers, the agent attempts to place them but has no knowledge of where they should go to produce a single continuous scaffold. (B) User-provided 2-by-6 rectangle template with correct crossover routing and a midseam that routes the scaffold in 1 continuous loop. The half-crossover vs full-crossover distinction and the midseam pattern were explained alongside this template. (C) After learning the midseam and half-crossover placement rules from (Figure 3, B), the agent generalizes to a 2-by-8 rectangle with correct routing, a single contiguous scaffold loop, without further instruction.

### 5.2 User Correction

The user provided a hand-designed 2-by-6 rectangle with correct dense crossover routing (Figure 3, B), along with an explanation of the placement rules. The key distinction is that at each helix pair boundary, the scaffold turns via a single half-crossover (left edge or right edge, determined by helix parity), while the interior is connected by double crossovers at every valid honeycomb position. This creates a “single midseam” pattern that produces one continuous scaffold oligo.

Additional errors in this phase included scaffold/staple strand assignment (the agent placed scaffold strands on the staple strand set and vice versa), which the user identified by inspecting the design in caDNAno’s strand routing view.

### 5.3 Generalization

After learning the midseam and half-crossover placement rules, the agent generalized to various 2-layer lattice dimensions without further instruction (Figure 3, C), each with a single contiguous scaffold loop. The first of these served as the end-to-end validation of the pipeline from design through molecular dynamics simulation.

## 6. Formalizing Cavity Design

With scaffold routing established for simple rectangles, we introduced a more complex design goal, a 2-layer rectangle with a rectangular cavity. The cavity is a gap in the interior of the structure along the helix length axis. We enforced the constraint that the total scaffold length could not exceed 8,064 nt (the p8064 scaffold). This introduced a second set of failure modes.

### 6.1 Initial Attempt

The agent’s first cavity attempt (Figure 4, A) correctly identified where strands should be absent, leaving a gap in the helix length axis. However, the knowledge acquired for simple rectangle routing (Section 5) did not transfer to the cavity case. The routing pattern changes at cavity boundaries. The single midseam must be interrupted, full crossover seams run down the left side, right side, and middle of the structure, and half-crossovers at the cavity edges define the scaffold turn points. The agent had no basis for placing these half-crossovers, and the bottom helix pair requires the midseam to be omitted entirely. The result was a fragmented scaffold with broken routing at the cavity boundaries.

**Figure 4:**
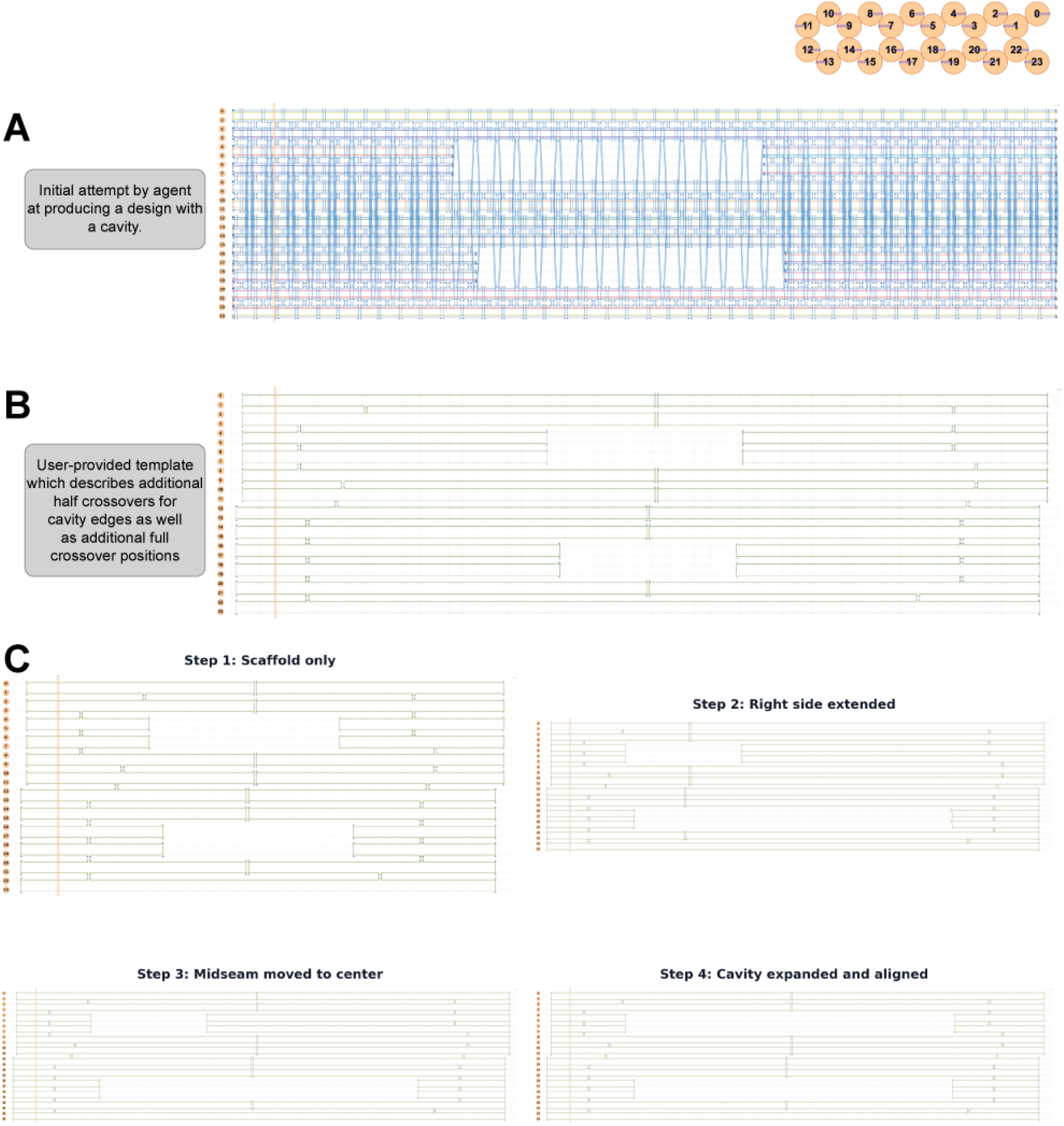
Formalizing cavity design. (A) Agent’s initial attempt at cavity routing. The agent correctly left a gap where strands should be absent, but could not place the half-crossovers needed to maintain scaffold continuity at the cavity boundaries. Knowledge from simple rectangle routing failed to transfer to the cavity case. (B) User-provided 2-by-12 template demonstrating correct cavity routing: full crossover seams on the left, right, and center of the structure, with the midseam omitted on the bottom helix pair, and half-crossovers defining the cavity edges. (C) After incorporating the user’s template, the agent developed a 4-step scaling process (scaffold only → extend right side → move midseam to center → expand and align cavity) to resize the structure and re-center the cavity at arbitrary dimensions.

### 6.2 User Correction

The user provided a 2-by-12 template with correct cavity routing (Figure 4, B), demonstrating the full crossover seam pattern and the half-crossover placement rules at cavity edges. This template, along with an explanation of how the routing differs from the simple rectangle case, was sufficient for the agent to incorporate the new pattern.

### 6.3 Scaling and End-to-End Validation

With cavity routing understood, the agent developed a 4-step process for resizing and re-aligning the cavity (Figure 4, C). After each step, the agent verified that the resulting JSON opened correctly in caDNAno without errors.

1. Remove staples (keep scaffold only).
2. Extend helix arrays and shift right-side crossovers.
3. Move midseam crossovers to center.
4. Expand cavity by moving boundary crossovers and re-aligning to the center.

Additional failures arose during scaling: the agent’s script left stray staple entries with broken crossover references, causing caDNAno to crash; the agent filled the cavity gap with continuous scaffold, destroying the cavity; and the scaling logic moved crossover positions independently per helix, creating skewed crossovers where both sides must share the same index. Each was resolved by user correction, and the agent did not repeat any of these error classes.

The resulting design was a 2-by-12 rectangular origami with an aligned cavity, 1 continuous scaffold loop (7,688 bp), scaled from the user’s 252 bp template to 420 bp helices.

## 7. Cumulative Capability and Targeted Edits

Each correction was retained across subsequent sessions; the agent did not repeat any resolved error class. The cumulative effect on agent capability was observed at each stage.

1. **After lattice geometry correction** (Section 4). All subsequent designs used correct 2-by-N grid dimensions.
2. **After scaffold routing correction** (Section 5). The agent adopted the user’s template as the basis for all designs, replacing from-scratch construction.
3. **After cavity orientation correction** (Section 6). All cavity designs placed the gap along the helix length axis.
4. **After scaling corrections** (Section 6). The agent developed a validated 4-step scaling process.

After correction of all failure modes, the agent operated autonomously on the following tasks without further human intervention.

1. **Extended the template to arbitrary lattice dimensions** (e.g., 2-by-16, 2-by-20) by adding column pairs to both sides of the base template, maintaining a single contiguous scaffold loop at each step.
2. **Designed to specification**. Given the constraint “20 nm × 40 nm cavity, centered, 6-helix padding, scaffold ≤ 8,064 bp,” the agent independently selected a 2-by-20 grid (40 helices, 252 bp), computed the cavity dimensions (8 columns, 117 bp gap), and built the complete design (7,828 bp scaffold, 1 oligo).

### Targeted Edits and Autonomous Pipeline Execution

On a 2-by-22 integrin cavity design, we observed that several scaffold crossovers were positioned too close to the cavity edge (3–5 bp from the boundary half-crossovers). We explicitly prompted the agent to move them, and it identified 5 inter-pair crossovers, relocated each to a valid lattice position away from the cavity, and recomputed centered cavity boundaries (Figure 5, A). The agent then read the caDNAno source for autoStaple and autoBreak and applied both to produce a stapled design.

**Figure 5:**
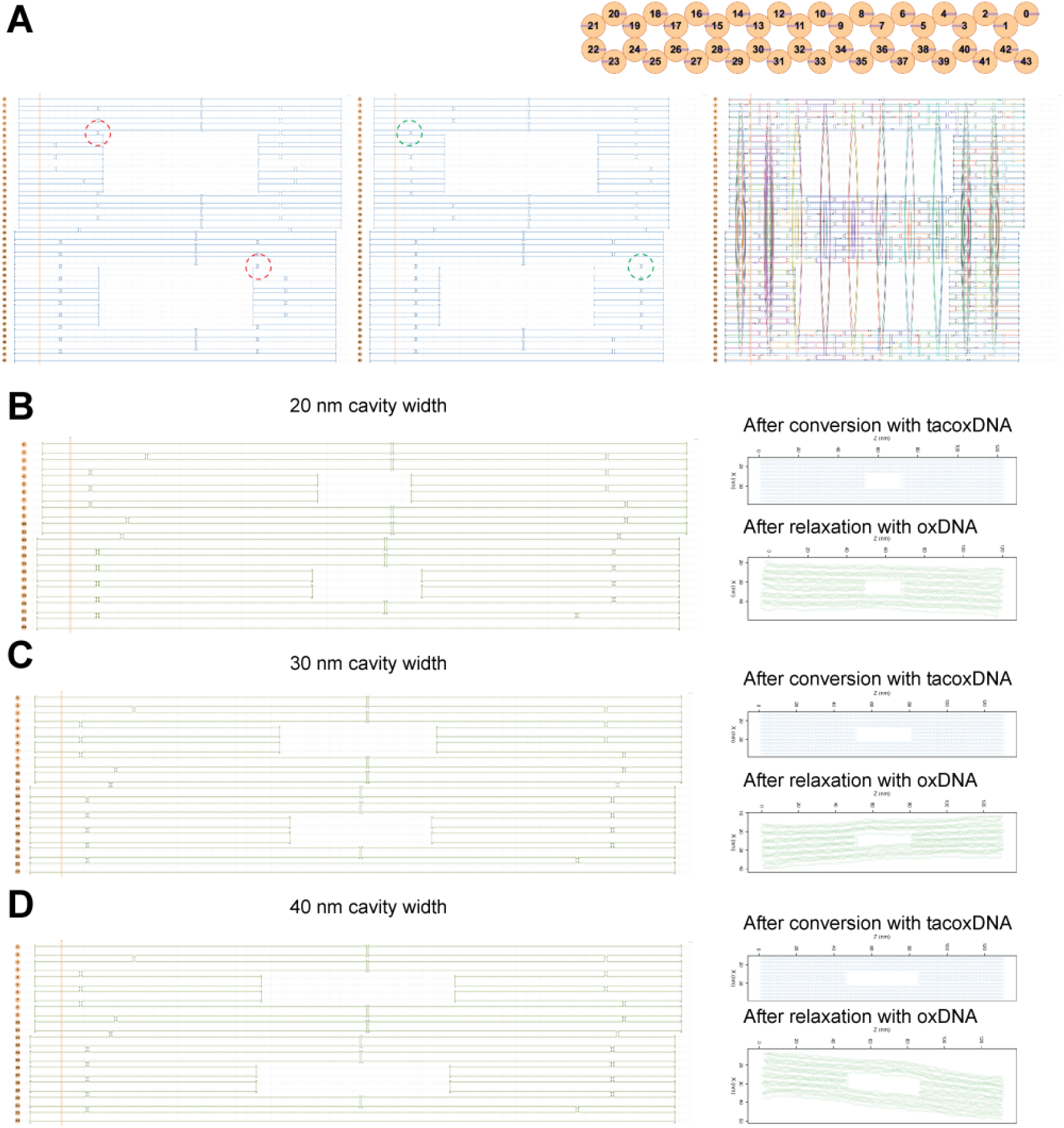
Autonomous targeted edits and end-to-end pipeline execution. (A) Crossover repositioning: the user prompted the agent to move scaffold crossovers away from the cavity edge, followed by autoStaple and autoBreak, producing a complete stapled design. (B–D) Parametric cavity variants at 20 nm, 30 nm, and 40 nm widths, each produced autonomously: caDNAno design (left), tacoxDNA-converted initial 3D structure (center), and oxDNA-relaxed structure (right). The agent computed cavity dimensions from physical specifications, built each design, stapled, converted, and ran molecular dynamics without user interaction.

### Parametric Cavity Variants

With the pipeline fully operational, the agent autonomously generated cavity variants at three widths (20 nm, 30 nm, and 40 nm) without user interaction (Figure 5, B–D). For each variant, the agent computed the cavity dimensions from physical specifications, built the caDNAno design, ran autoStaple and autoBreak, converted to oxDNA format via tacoxDNA, and submitted molecular dynamics relaxation on a GPU computing cluster. The resulting oxDNA structures maintained the cavity shape after relaxation, confirming structural validity.

Each of these tasks required the agent to apply domain knowledge transferred in previous sessions: crossover lattice positions, parity-dependent scaffold direction, cavity boundary half-crossover placement, and honeycomb valid offsets. None were re-taught. The corrections from prior sessions accumulated as reusable code (verifier checks in `cadnano_verifier.py`, pipeline functions in `cavity_variant_sweep.py`, lessons in `tasks/lessons.md`) that the agent applied automatically in new design contexts.

## 8. Discussion

The agent cannot design DNA origami independently. However, the failure modes documented in this work each encode domain knowledge with three properties.

1. **Invisible in source code**. Rules such as the grid-to-layer mapping, the midseam routing pattern, and the crossover directionality conventions at cavity boundaries cannot be deduced from caDNAno’s codebase. They require explicit transfer from an experienced designer.
2. **Retained after single correction**. Once corrected, the agent does not repeat that error class. The correction is encoded in verifier functions, lessons files, and pipeline logic that persist across sessions.
3. **Distributable**. The verifier (`cadnano_verifier.py`) checks all documented failure modes automatically. The failure catalog (`failure_analysis.md`) documents each with symptom, root cause, fix, and verifier check. A new user or agent can apply these checks without repeating the original debugging.

The agent functions not as a designer but as a substrate for accumulating and distributing design knowledge (Figure 1, C). Each user-agent interaction produces reusable verification and pipeline tools in addition to the design itself. Table 1 lists the artifacts from this work that are distributable to any DNA origami researcher working with caDNAno.

**Table 1:** Distributable artifacts from this work.

| Artifact | Purpose | Reusability |
| --- | --- | --- |
| `cadnano_verifier.py` | Pre-flight design validation | Run on any caDNAno JSON before committing to synthesis |
| `failure_analysis.md` | Documented failure modes | Read before starting AI-assisted design; avoid known pitfalls |
| `cavity_variant_sweep.py` | Parametric cavity pipeline | Change gap size/width → design recomputes automatically |
| Template extension functions | Add columns to existing designs | Extend any 2-by-N template to 2-by-(N+2) with a single contiguous scaffold loop |
| Crossover move pattern | Crossover directionality template | Apply to any targeted crossover edit on any honeycomb design |
| Renumbering procedure | JSON helix renumbering | Essential for any template extension; ensures parity correctness |

The pipeline from the user’s template to verified design to oxDNA simulation is end-to-end, requiring no manual caDNAno GUI interaction. For example, a researcher can describe a cavity design in terms of physical dimensions (nm × nm, scaffold length, padding) and receive a validated JSON, stapled design, and oxDNA files ready for submission. Integration with cluster scheduling can extend this further, including 3D analysis functions that automatically infer and flag structural properties from molecular dynamics simulation trajectories.

The transition from demonstration to knowledge transfer occurred when the agent began generating verification tools from its own failure modes. The verifier was not designed top-down. It emerged from specific debugging sessions, each producing a check that prevents the corresponding failure for any subsequent user.

Returning to the question posed in the introduction, whether a coding agent could serve as a unified front end for DNA origami design, the answer from this initial investigation is qualified. The agent cannot replace domain expertise, but it can serve as a mechanism for capturing that expertise in executable form. The design rules formalized in this work began as corrections delivered in natural language and ended as automated scripts, verifier checks, and documented constraints that any agent or user can apply. This suggests a model in which each new design challenge, once resolved through agent-designer interaction, permanently reduces the friction of that task class for all subsequent users. Whether this model scales to the full diversity of DNA origami structures, like curved, wireframe, or complex 3D structures, remains to be demonstrated, but this work sets a starting point for investigating coding agents as a consolidating control surface for DNA origami design, one that folds in and combines existing and emerging design methodologies into a streamlined pipeline.

## Appendix A Agent Operations Library

Examples of natural language prompt instructions and the corresponding agent output for targeted modifications on a 6-helix design. Each row shows the before state (left) and after state (right) for a single operation. Operations include crossover creation, crossover movement, crossover deletion, strand extension and shrinking, insertion and deletion placement, strand splitting, automatic staple breaking, and bulk crossover operations.

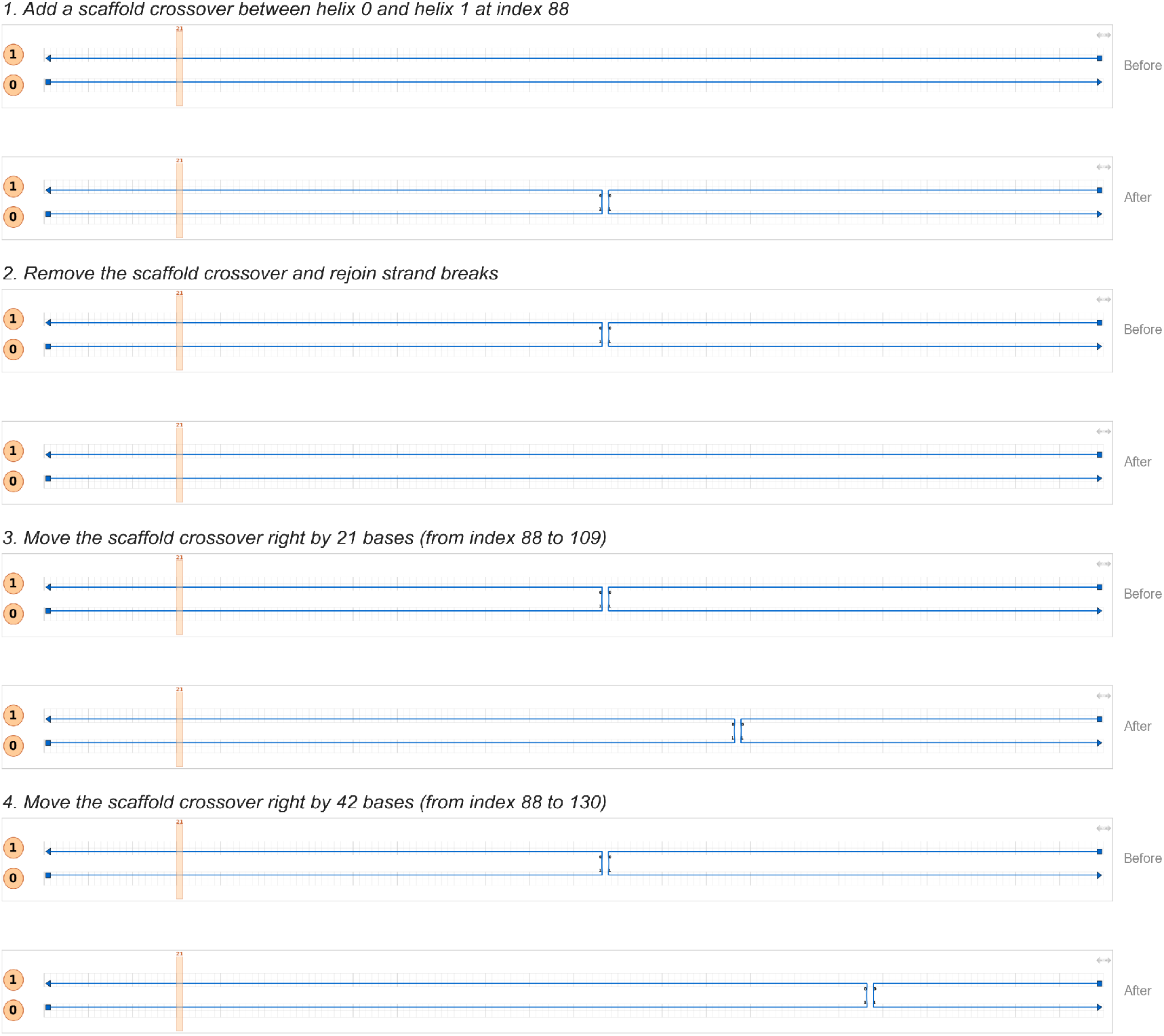
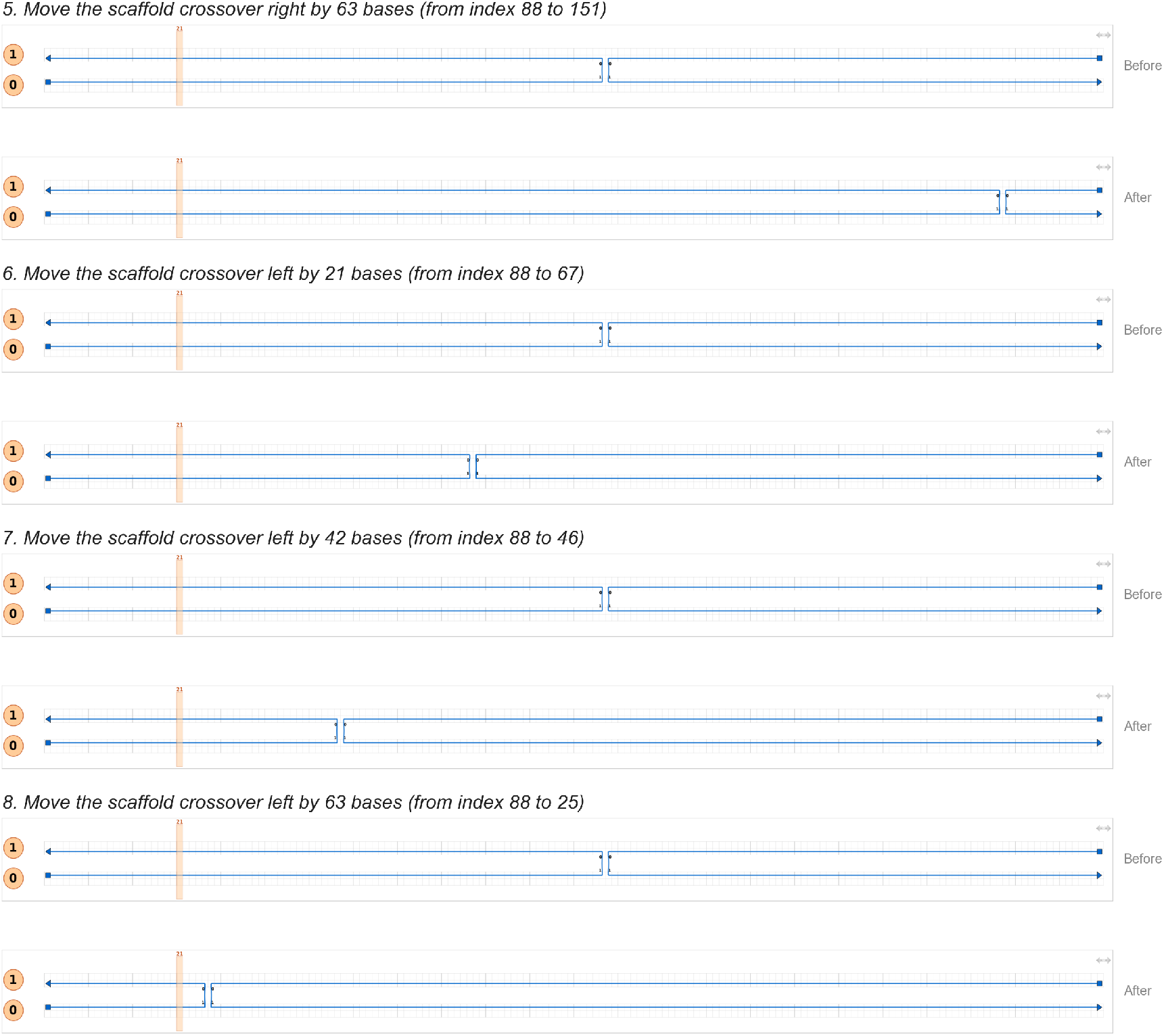
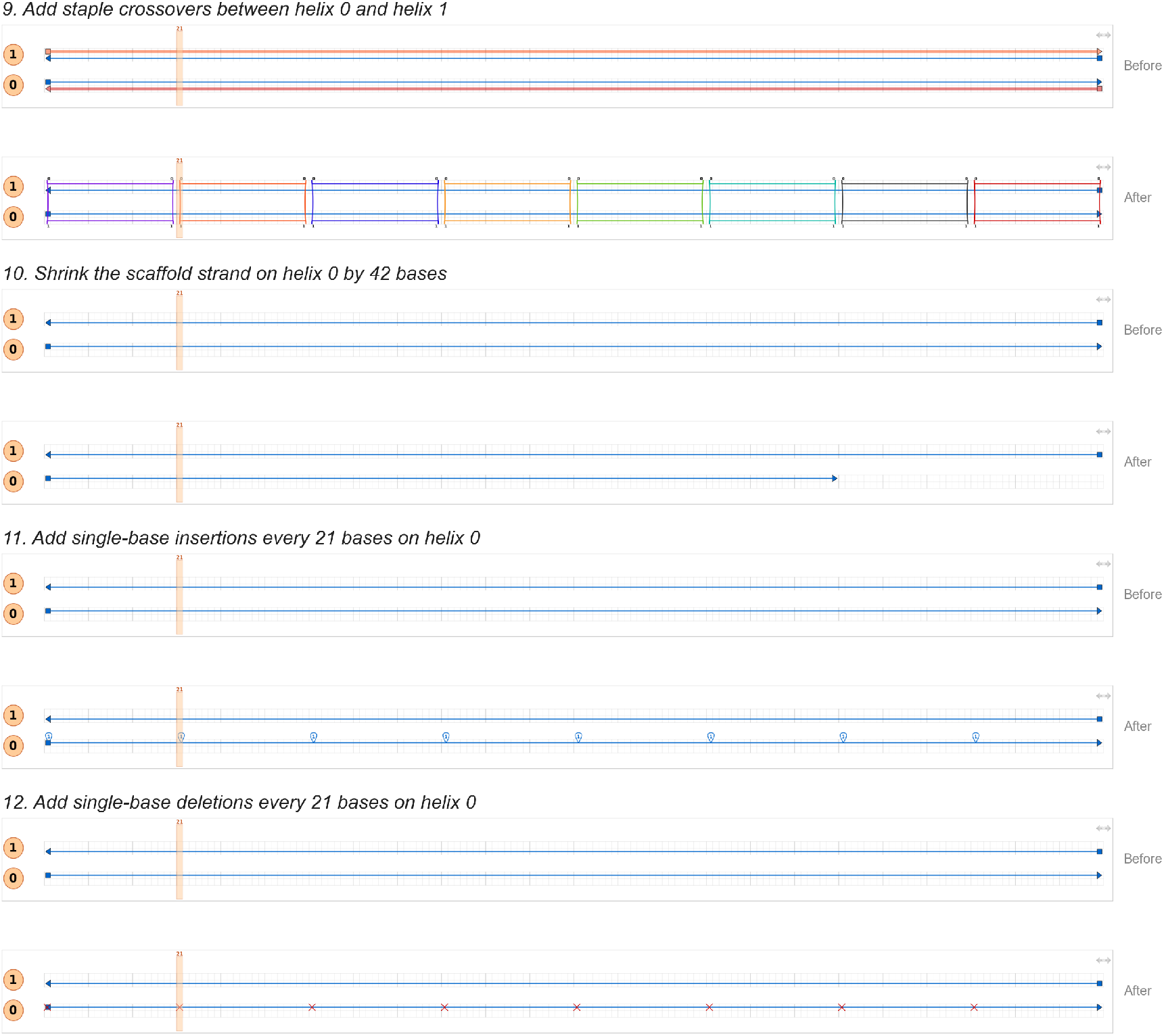

## Notes

### Competing Interest Statement

The authors have declared no competing interest.

## References

(1) Majikes, J. M.; Liddle, J. A. DNA Origami Design: A How-to Tutorial. Journal of Research of the National Institute of Standards and Technology 2021, 126, 126001.

(2) Aksel, T.; Navarro, E. J.; Fong, N.; Douglas, S. M. Design Principles for Accurate Folding of DNA Origami. Proceedings of the National Academy of Sciences 2024, 121 (48).

(3) Douglas, S. M.; Marblestone, A. H.; Teerapittayanon, S.; Vazquez, A.; Church, G. M.; Shih, W. M. Rapid Prototyping of 3d DNA-Origami Shapes with Cadnano. Nucleic Acids Research 2009, 37 (15), 5001–5006.

(4) Fu, D.; Narayanan, R. P.; Prasad, A.; Zhang, F.; Williams, D.; Schreck, J. S.; Yan, H.; Reif, J. Automated Design of 3d DNA Origami with Non-Rasterized 2d Curvature. Science Advances 2022, 8 (51).

(5) Veneziano, R.; Ratanalert, S.; Zhang, K.; Zhang, F.; Yan, H.; Chiu, W.; Bathe, M. Designer Nanoscale DNA Assemblies Programmed from the Top down. Science 2016, 352 (6293), 1534–1534.

(6) Benson, E.; Mohammed, A.; Gardell, J.; Masich, S.; Czeizler, E.; Orponen, P.; Högberg, B. DNA Rendering of Polyhedral Meshes at the Nanoscale. Nature 2015, 523 (7561), 441–444.

(7) Jun, H.; Zhang, F.; Shepherd, T.; Ratanalert, S.; Qi, X.; Yan, H.; Bathe, M. Autonomously Designed Free-Form 2d DNA Origami. Science Advances 2019, 5 (1).

(8) Jun, H.; Wang, X.; Bricker, W. P.; Bathe, M. Automated Sequence Design of 2d Wireframe DNA Origami with Honeycomb Edges. Nature Communications 2019, 10 (1).

(9) Babatunde, B.; Arias, D. S.; Cagan, J.; Taylor, R. E. Generating DNA Origami Nanostructures Through Shape Annealing. Applied Sciences 2021, 11 (7), 2950.

(10) Levy, N.; Schabanel, N. Ensnano: A 3d Modeling Software for DNA Nanostructures. DNA 27, Leibniz International Proceedings in Informatics 2021, 205, 5:1–5:23.

(11) Rothemund, P. W. K. Folding DNA to Create Nanoscale Shapes and Patterns. Nature 2006, 440 (7082), 297–302.

(12) Douglas, S. M.; Dietz, H.; Liedl, T.; Högberg, B.; Graf, F.; Shih, W. M. Self-Assembly of DNA into Nanoscale Three-Dimensional Shapes. Nature 2009, 459 (7245), 414–418.

(13) Ke, Y.; Douglas, S. M.; Liu, M.; Sharma, J.; Cheng, A.; Leung, A.; Liu, Y.; Shih, W. M.; Yan, H. Multilayer DNA Origami Packed on a Square Lattice. Journal of the American Chemical Society 2009, 131 (43), 15903–15908.

(14) Dietz, H.; Douglas, S. M.; Shih, W. M. Folding DNA into Twisted and Curved Nanoscale Shapes. Science 2009, 325 (5941), 725–730.

(15) Ke, Y.; Voigt, N. V.; Gothelf, K. V.; Shih, W. M. Multilayer DNA Origami Packed on Hexagonal and Hybrid Lattices. Journal of the American Chemical Society 2012, 134 (3), 1770–1774.

(16) He, J.; Treude, C.; Lo, D. LLM-Based Multi-Agent Systems for Software Engineering: Literature Review, Vision, And the Road Ahead. ACM Transactions on Software Engineering and Methodology 2025, 34 (5), 1–30.

(17) Dong, Y.; Jiang, X.; Qian, J.; Wang, T.; Zhang, K.; Jin, Z.; Li, G. A Survey on Code Generation with LLM-Based Agents. arXiv preprint 2508.00083 2025.

(18) Suma, A.; Poppleton, E.; Matthies, M.; Šulc, P.; Romano, F.; Louis, A. A.; Doye, J. P. K.; Micheletti, C.; Rovigatti, L. Tacoxdna: A User-Friendly Web Server for Simulations of Complex DNA Structures, From Single Strands to Origami. Journal of Computational Chemistry 2019, 40 (29), 2586–2595.

(19) Ouldridge, T. E.; Louis, A. A.; Doye, J. P. K. Structural, Mechanical, And Thermodynamic Properties of a Coarse-Grained DNA Model. The Journal of Chemical Physics 2011, 134 (8), 85101.

(20) Šulc, P.; Romano, F.; Ouldridge, T. E.; Rovigatti, L.; Doye, J. P. K.; Louis, A. A. Sequence-Dependent Thermodynamics of a Coarse-Grained DNA Model. The Journal of Chemical Physics 2012, 137 (13), 135101.

(21) Snodin, B. E. K.; Randisi, F.; Mosayebi, M.; Šulc, P.; Schreck, J. S.; Romano, F.; Ouldridge, T. E.; Tsukanov, R.; Nir, E.; Louis, A. A.; Doye, J. P. K. Introducing Improved Structural Properties and Salt Dependence into a Coarse-Grained Model of DNA. The Journal of Chemical Physics 2015, 142 (23), 234901.

(22) Rovigatti, L.; Šulc, P.; Reguly, I. Z.; Romano, F. A Comparison between Parallelization Approaches in Molecular Dynamics Simulations on GPUs. Journal of Computational Chemistry 2015, 36 (1).

(23) Guo, D.; Yang, D.; Zhang, H.; Song, J.; Wang, P.; Zhu, Q.; Xu, R.; Zhang, R.; Ma, S. Deepseek-R1: Incentivizing Reasoning Capability in LLMs via Reinforcement Learning. Nature 2025, 645 (8081), 633–638.

(24) Wang, X.; Chen, Y.; Yuan, L.; Zhang, Y.; Li, Y.; Peng, H.; Ji, H. Executable Code Actions Elicit Better LLM Agents. In Proceedings of the 41st International Conference on Machine Learning (ICML); 2024.

